# IC-VAE: A Novel Deep Learning Framework for Interpreting Multiplexed Tissue Imaging Data

**DOI:** 10.1101/2023.11.06.565771

**Authors:** Huy Nguyen, Hy Vuong, Thao Truong, Son Pham

## Abstract

Interpreting protein expression in multiplexed tissue imaging data presents a significant challenge due to the high dimensionality of the resulting images, the variety of intracellular structures, cell shapes resulting from 2-D tissue sectioning, and the presence of technological noise and imaging artifacts. Here, we introduce the Information-Controlled Variational Autoencoder (IC-VAE), a deep generative model designed to tackle this challenge. The contribution of IC-VAE to the VAE framework is the ability to control the shared information among latent subspaces. We use IC-VAE to factorize each cell’s image into its true protein expression, various cellular components, and background noise, while controlling the shared information among some of these components. Compared with other normalization methods, this approach leads to superior results in downstream analysis, such as analyzing the expression of biomarkers, classification for cell types, or visualizing cell clusters using t-SNE/UMAP techniques.

## 1 Introduction

Recent advances in highly multiplexed tissue imaging technologies, such as multiplexed immunofluorescence and imaging mass spectrometry [1] have enabled the simultaneous measurement of various proteins at single-cell resolution. Central to the analysis of highly multiplexed imaging data is the accurate identification of individual cells and the quantification of protein expression levels within them. Most current workflows [2] follow the following steps:

- Perform cell segmentation on the nucleus staining image and dilate the segmentation to obtain cell membrane. (As membrane staining is less reliable than nucleus staining, and current cell segmentation algorithms work better with nucleus segmentation).
- Quantify protein expression by summing the intensity of each protein for pixels within the cell boundaries.
- Normalize the protein expression of each cell by the corresponding cell area.

This workflow faces several limitations. First, the dilating procedure assumes the symmetry property of cells in the tissue section, which is often violated. Second, the tissue section cuts cells at different angles, resulting in different cell compartments (e.g., membrane, nucleus, cytoplasm, etc) in the image. While the protein signal is captured differently in cell compartments (e.g., proteins on the membrane will more likely be captured in the membrane compartment), normalizing the total expression by the cell area implicitly admits proteins can appear in any compartment in a cell at the same rate. Additionally, in areas where cells are densely located, signals from neighboring cells create a noisy background and should be eliminated.

We here propose a model to accurately quantify protein expression by using variational autoencoder (VAE) to reconstruct the original images of each cell using 3 components: the protein expression, the compartments of each cell, and background noise. We build a neural network to calculate and minimize the mutual information among the components. Through multiple benchmarks, we show that our new approach is more accurate than MCMICRO [2], the current state-of-the-art approach for quantifying protein expression on highly multiplexed tissue images.

## 2 Methods

Given an image A of dimensions *h×w*, which captures the signal corresponding to protein distribution within a cell, we observe that the signal intensity varies across the image. This variation arises because different cellular compartments possess varying quantities of protein molecules, a heterogeneity intrinsic to the biological nature of the cell. Let r represent the number of distinct cellular compartments (e.g., nucleus, cytoplasm, membrane, etc.) within the cell, and let *s* denote the aggregate number of protein molecules present in the cell. Then, we can model the distribution of protein signals in the image as follows:

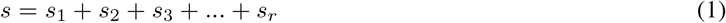

where *s*_*i*_ is the number of protein molecules in the *i*-th cell compartment, *s*_*i*_ *≥* 0.

Furthermore, the image *A* captures a two-dimensional cross-section of a three-dimensional cell, which leads to the variability of protein signal within the same cell compartment. Let us define a *h × w* matrix

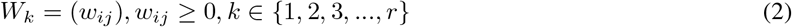

representing the extent to which the position *i, j* on the sectioning plane of the *k*-th cell compartment influences the protein signal magnitude.

From (1) and (2), we can interpret the image *A* as:

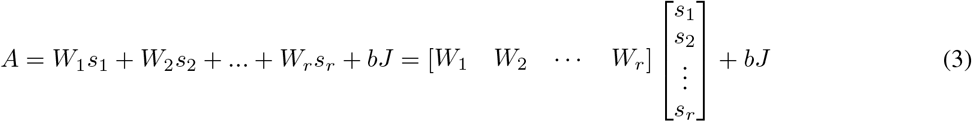

where *b* represents the background noise and imaging artifacts, *J* is the matrix of ones with the same dimension as *A* (and each *W*_*k*_).

In a highly multiplexed tissue imaging dataset, scientists measure the signal of multiple proteins. Let *c* be the number of proteins. Assuming the effect of cell compartments on measuring protein signals is identical among proteins. Based on (3), we have *c* equations:

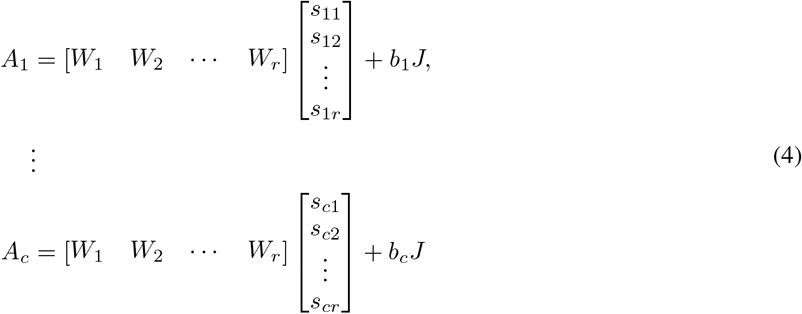

Let *t* = *h·w*. The images *A*_1_, *A*_2_, …, *A*_*c*_ can be represented as a matrix *V* of size *t×c* where each column contains *t* pixel values of one of the *c* images in the set, flattened into a vector. Therefore, equation (4) is equivalent to:

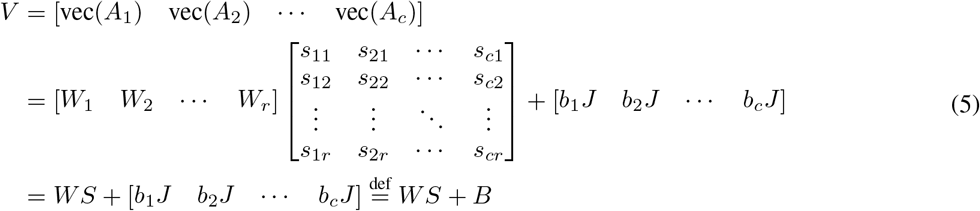

where *S* is a *r×c* matrix, *s*_*ij*_ is the number of molecules of the *j*-th protein at the *i*-th cell compartment, *W* is a *t × r* matrix where each column contains the flattened values of *W*_*i*_, with *J* being a vector of length *t* with all elements equal to 1. *B* is a *t × c* matrix, each *i*-th columns contains *t* flattened values of *b*_*i*_*J*.

We are interested in *S*. By summing elements within the columns of *S*, we obtain a *c*-dimensional vector 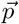 containing the total number of molecules of each protein:

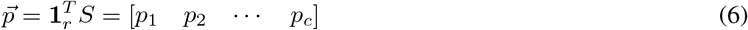

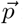 is the protein expression level that we want to quantify given *V* . As a multiplexed tissue imaging dataset contains multiple cells, by solving (5) iteratively over all the cells, we obtain the protein expression matrix:

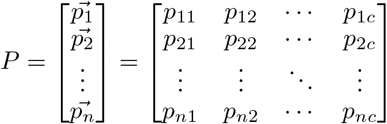

where *n* is the number of cells, *c* is the number of protein, *p*_*ij*_ *≥*0, *∀* 0≤ *i*0≤ *n*, ≤ *j*≤ *c*.

To construct *P*, we introduce a deep learning framework called Information-Controlled Variational Autoencoder (IC-VAE). The principal steps are as follows:

First, we design a VAE-based neural network that encodes the protein signal images of each cell to a latent distribution. From this latent distribution, we sample three latent subspaces:

- The protein expression latent subspace *Z*_*expression*_.
- The cell compartments latent subspace *Z*_*compartments*_.
- The background noise latent subspace *Z*_*background*_.

We then decode the latent representations to obtain *W*_*decoded*_, *S*_*decoded*_, *B*_*decoded*_ and reconstruct the original image as *W*_*decoded*_*S*_*decoded*_ + *B*_*decoded*_. The decoding process includes normalization steps: each compartment within *W*_*decoded*_ is normalized to control for compartment-specific effects, and normalization across cells is applied to ensure comparability of the protein expression profiles between different cells. Moreover, to maintain the distinctiveness of each latent subspace, we incorporate a specialized neural network to calculate and minimize the Mutual Information (MI) among them. This step ensures that the subspaces contain non-redundant, independent features relevant to their respective roles in reconstructing the protein signal images.

### 2.1 IC-VAE Algorithm

#### Inputs

We describe how to construct the input for our method - IC-VAE, given a multiplexed tissue imaging dataset *D* containing *c* protein signal images of size *h×w*. Since we aim to quantify the protein expression level for each cell in the dataset, we first need to identify individual cells on the image. Our method does not rely on membrane segmentation of the cell. Therefore, readers can apply state-of-the-art cell segmentation methods on the nucleus staining channel of the dataset. In this work, we use cellpose v2.2 [3], model *cyto*2, with default parameters to obtain the nucleus segmentation mask.

The nucleus segmentation mask *A* = (*a*_*i,j*_) is a *h× w* matrix, where *a*_*i,j*_ = 0 indicates no cell detected at the pixel (*i, j*), *a*_*i,j*_ = *k* indicates the pixel (*i, j*) belongs to cell *k*^*th*^.

Using the nucleus segmentation mask *A*, for each cell, we extract an equal size *m× m* image for every protein, such that the center of the cell is at the position (*m//*2, *m//*2) of the image. Then, we normalize the images to the range [0, 1] and construct a tensor *T* of size *c × m × m* representing one cell. The set *𝒳* = *{T*_1_, *T*_2_, …, *T*_*n*_*}*, with *n* is the number of cells in *D*, is the input for our model.

For the training convention, we split *𝒳* into two sets with the ratio 9 : 1: the training set *𝒳* _*training*_ and the validation set *𝒳* _*validation*_. During training, we feed *X*_*training*_ to IC-VAE in batches of size *N*. Following the best practices, we shuffle *𝒳* _*training*_ for each epoch and use the *𝒳* _*validation*_ to validate the model after every epoch to avoid overfitting.

Notice that the choice of *m* varies between datasets. We want to select *m* large enough so that the biggest cell in *D* completely lies within the sub-image and is small enough for efficient training performance. In practice, we usually select *m* = 128.

#### Hyperparameters

Our method requires one hyperparameter *r* - which represents the number of cell compartments. We choose *r* such that *r << nc*. In practice, we choose *r* = 4 - under the intuition that *r* = 4 represents the cell nucleus, the cell cytoplasm, the cell membrane, and the rest.

#### Model architecture

Here, we outline the general architecture of IC-VAE. (Figure 1).

**Figure 1:**
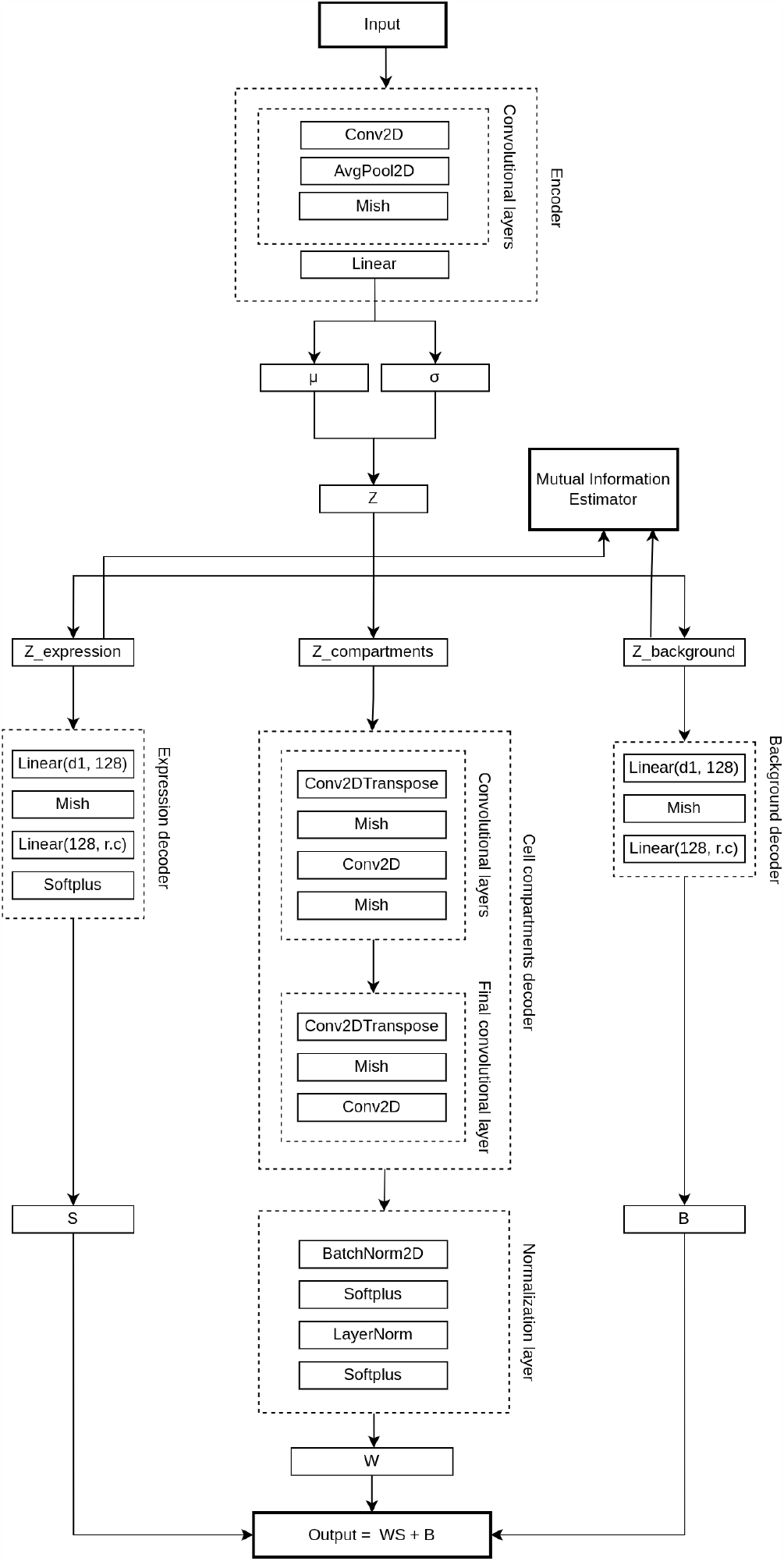
Information-Controlled Variational Auto-Encoder (VAE) based architecture for image analysis. The encoder consists of convolutional layers that process the input image and map it to a latent space representation, *Z*. The latent representation is then used by three separate decoders: the background decoder, the cell compartments decoder, and the expression decoder. Each decoder consists of convolutional layers and reconstructs different aspects of the image. The final output is normalized before being outputted. The Mutual Information Estimator component is used to control the shared information between *Z*_*expression*_ and *Z*_*background*_.

##### 2.1.1 The encoder

The encoder receives a tensor of size *N×c×m×m* as input. The input is passed through a series of convolutional layers, followed by a linear layer to simultaneously obtain the *μ* and *σ* of the latent distribution. Using the reparameterization trick [4], we obtain the *d*-dimensional latent space *Z* = *μ* + *ϵ·σ, ϵ∈ 𝒩* (0, 1). Z is a *N× d* matrix. Each row is a latent vector representation of the corresponding cell. Next, we split *Z* into three sub-matrices:

- A *N × d*_1_ matrix *Z*_*expression*_ represents the *d*_1_-dimensional protein expression latent subspace.
- A *N × d*_2_ matrix *Z*_*compartment*_ represents the *d*_2_-dimensional cell compartments latent subspace.
- A *N × d*_3_ matrix *Z*_*background*_ represents the *d*_3_-dimensional background noise latent subspace.

where *d* = *d*_1_ + *d*_2_ + *d*_3_. Here, the choice of *d*_1_, *d*_2_, and *d*_3_ will influence the resulting protein expression matrix. In practice, we usually choose 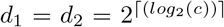 and 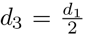 . Also, the number of convolutional layers and the number of in/out channels per convolutional layer varies between datasets. Empirical evidence from our experiments across several datasets suggests that a shallower encoder architecture tends to yield more desirable results in this specific application than a deeper one.

##### 2.1.2 The decoders

Instead of having a single decoder, we design three decoders with different architectures to obtain the desired outputs:

#### Expression decoder

The expression decoder transforms *Z*_*expression*_ to a *N×rc* matrix through a series of linear layers ending with the softplus activation function to ensure the non-negativity constraint. We reshape this matrix to a tensor *S*_*decoded*_ of size *N × r × c*, each *r × c* matrix from this tensor is *S* from equation (5) of the corresponding cell.

#### Cell compartments decoder

The cell compartments decoder transforms *Z*_*compartments*_ to a tensor of size *N×r×m×m* through a series of convolutional layers, each sub-tensor *r×m×m* is the matrix *W* from equation (5). We further pass *W* through a normalization layer to constrain the scale of *W*, which also constrains the scale of *S*. The cell compartments decoder also ends with a softplus activation function to ensure the non-negativity constraint and produce the final output *W*_*decoded*_.

#### Background decoder

The background decoder transforms *Z*_*background*_ to a tensor *B*_*decoded*_ of size *N × c* through a series of linear layers with no activation function layer at the end to allow the output to be at any scale. Besides using the output tensor to obtain *B* from equation (5), we also want to use this tensor as a bias for later reconstruction. One could represent the output of this layer as *B*_*decoded*_ = *B* + *Bias*.

#### Mutual Information

We use MINE algorithm [5] to calculate the mutual information between *Z*_*expression*_ and *Z*_*background*_ with the goal of minimizing the shared information between them. Details about the MINE algorithm are described in the Supplementary Materials.

#### Reconstruction

After having *S*_*decoded*_, *W*_*decoded*_, and *B*_*decoded*_, we apply the Einstein summation with the convention *Ncm, Nmhw → Nchw* on *S*_*decoded*_ and *W*_*decoded*_ to obtain the output tensor *O* of size *N × c × m × m*. Then, we broadcast the tensor *B*_*decoded*_ to a tensor of size *N×c×m×m* and add this tensor to *O* to obtain the reconstructed images.

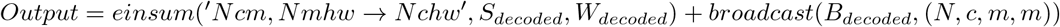

#### Loss function

Our IC-VAE loss comprises the ELBO loss in variational inference and mutual information loss. The ELBO loss comprises the expectation of log-likelihood and the weighted KL divergence. The expectation of log-likelihood corresponds to the difference between the input and the reconstructed images from the latent representation. Weighted KL-divergence loss corresponds to the distance between the posterior and the chosen prior. This helps regularize the latent space.

We denote the prior of the latent space as *p*(*z*), the likelihood of the sample given the latent representation as *p*(*x*|*z*) computed from a decoder *D*(*z*), the posterior of the latent space given the sample as *p*(*z*|*x*)

To make it feasible, we approximate the posterior *p*(*z*|*x*) as *q*(*z*|*x*) using an encoder *E*(*x*) = (*E*_*μ*_(*x*), *E*_*σ*_(*x*)) We chose our prior *p*(*z*) to be a normal distribution *z ∼ 𝒩* (0, 1), our likelihood to be *x*|*z ∼ 𝒩* (*D*(*z*), *I*) and our posterior *z*|*x ∼ 𝒩* (*E*_*μ*_(*x*), diag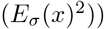

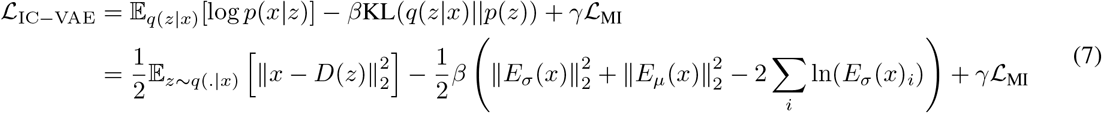

We denote the parameterized mutual information estimator as *MI*(*z*_exp_, *z*_bg_).

We define the mutual information loss, adopted from [6], the Jensen-Shannon Divergence as,

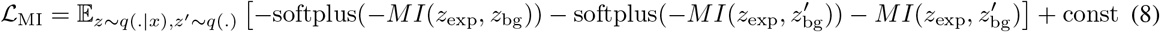

For each mini-batch, we first optimize ℒ_IC− VAE_ while fixing the parameters of ℒ _*MI*_, then we optimize _MI_ while fixing the parameters of the decoder *D* and the encoder *E*.

## 3 Benchmarks

For our experiment, we compare IC-VAE against MCMICRO. To assess the efficiency of each method, we use the protein expression matrix that both methods generated as input for evaluating some common downstream analysis tasks, including analyzing the expression level of a protein, clustering cells for cell typing, and presenting the cell clusters on UMAP [7] plots.

### 3.1 Dataset description

We use the TONSIL-1 40X image of the first batch from a human tonsil cancer t-CyCIF dataset [8]. The image is a whole-slide 2-D image of size 16, 843*×*16, 125 pixels. It includes 44 different staining channels for cell nuclei, background control, and protein biomarkers. Imaging parameters and preprocessing methodology (e.g., background and shading correction, stitching, and registration) are described in [8].

### 3.2 Data pre-processing

#### 3.2.1 Cell segmentation

We select the first nucleus staining channel to perform cell segmentation. We use cellpose v2.2 [3] with *model* = *cyto*2 and *diameter* = *None*. Other parameters are kept as default. The resulting segmentation masks are later used as input for MIMICRO and IC-VAE.

#### 3.2.2 Cell-type labeling

To evaluate the clustering and dimensional reduction results, BioTuring’s pathologist manually annotates the cell type labels for three different sub-regions. The detailed description is presented in the supplementary materials.

#### 3.2.3 MCMICRO settings

We run the MCMICRO pipeline - revision 01c11e5615 (https://github.com/labsyspharm/mcmicro) using Nextflow version 23.04.4. For each dataset, we use the OME-TIFF whole-slide 2-D image and the cell segmentation mask from 3.2.1 as the input. Since we already have the processed image and cell segmentation, we configure the pipeline to start at the *quantification* step and stop at the *downstream* step. Other parameters are kept as default. The pipeline produces a *CSV* file containing the protein expression matrix.

#### 3.2.4 IC-VAE settings

We set the number of cell compartments *r* = 4. The mutual information weight *γ* is initialized with the value 1e*−* 5 and increased by 1e*−* 5 for every epoch until it reaches 1e*−* 3. The KL weight *β* is initialized with 1e*−* 5 and increased by 1e*−* 5 for every epoch until it reaches 1e*−* 4. We select the dimension for the latent subspaces as *d*_1_ = 32, *d*_2_ = 32, *d*_3_ = 16. We train the model with *batch−*_*size* = 128 and *n*_*epochs* = 100. If the validation loss is not increased after 10 epochs, we stop the training and use the best checkpoint to evaluate the entire input *X*.

#### 3.2.5 Downstream analysis

We use standard scanpy [9] pipeline for the downstream analysis benchmarks, including:

- PCA: For MCMICRO, we use PCA with default parameters. For IC-VAE, as described in section 2, the encoder produces an expression latent space. We utilize this latent space as a compressed representation of the data (similar to PCA).
- k-NN with *n*_*neighbors* = 30. Other parameters are kept as default.
- UMAP with *min*_*dist* = 0.1. Other parameters are kept as default.
- Leiden clustering with *resolution* = 1.0. Other parameters are kept as default.

Then, we perform the clustering and UMAP on the labeled cells from selected regions only. We combine three different regions to evaluate the ability to overcome the technical variants among cells from different field-of-views between MCMICRO and ICVAE.

### 3.3 Experimental results

#### 3.3.1 Analyzing the expression level of a protein

To evaluate the ability to overcome the background noise and imaging artifacts of the MCMICRO and IC-VAE, we select a region from the evaluated dataset where the signal of *CD*3*D* is highly affected by the aforementioned factors. As shown in Figure 2a, the right-hand side part of this region contains the background signal of *CD*3*D*, while the left side does not. We create a label called “Background region” and annotate the cells on the left side as “no background” and the cells on the right side as “have background” (Figure 2b). We also create a label called “CD3D expression” and annotate the cells in this region as *CD*3*D*^+^ - if the cell has strong *CD*3*D* signals surrounding its nucleus - otherwise, the cell is annotated as *CD*3*D*^−^ (Figure 2c).

**Figure 2:**
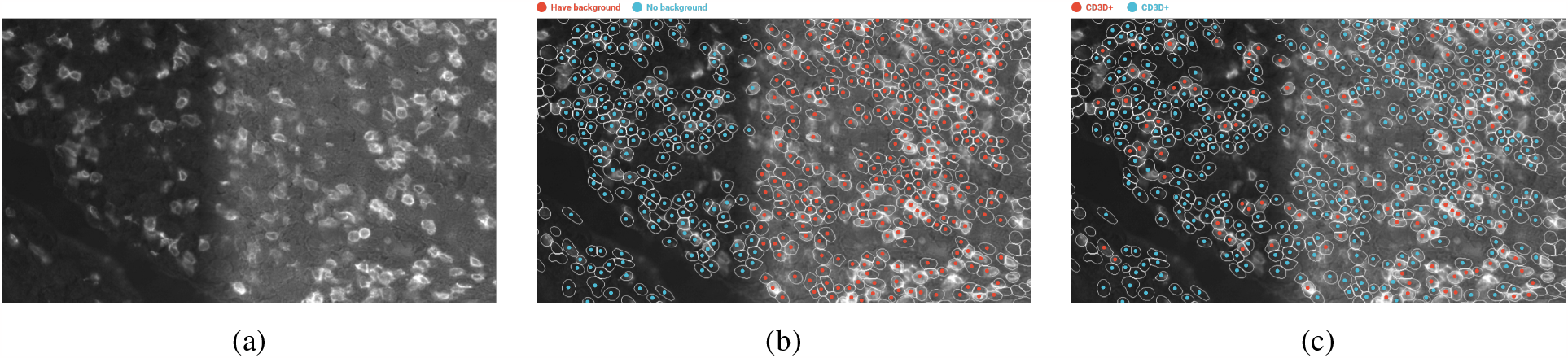
The region is highly affected by imaging artifacts. (a) The fluorescent image of *CD*3*D*. (b) Cells labeled by artifacts region. The blue dots stand for cells in the region without the background signal. The red dots stand for cells in the region with the background signal. (c) Cells labeled by protein expression. The blue dots stand for *CD*3*D*^−^ cells and the red dots stand for *CD*3*D*^+^ cells. The label is manually annotated with the same cell typing procedure described in the Supplementary Material.

Figure 3a indicates the background signal heavily influences the expression of *CD*3*D* quantified by MCMICRO. From the MCMICRO result, the expression of *CD*3*D* in *CD*3*D*^−^ cell population inside of the background noise region is even higher than the expression of *CD*3*D* in *CD*3*D*^+^ cell population outside of the background noise region (Figure 4a). On the other hand, in the result of IC-VAE model, the influence of the background signal on the *CD*3*D* expression is reduced (Figure 3b), the expression of *CD*3*D* in *CD*3*D*^−^ inside of the background noise region is lower than the expression of *CD*3*D* in *CD*3*D*^+^ cell population outside of the background noise region (Figure 4b).

**Figure 3:**
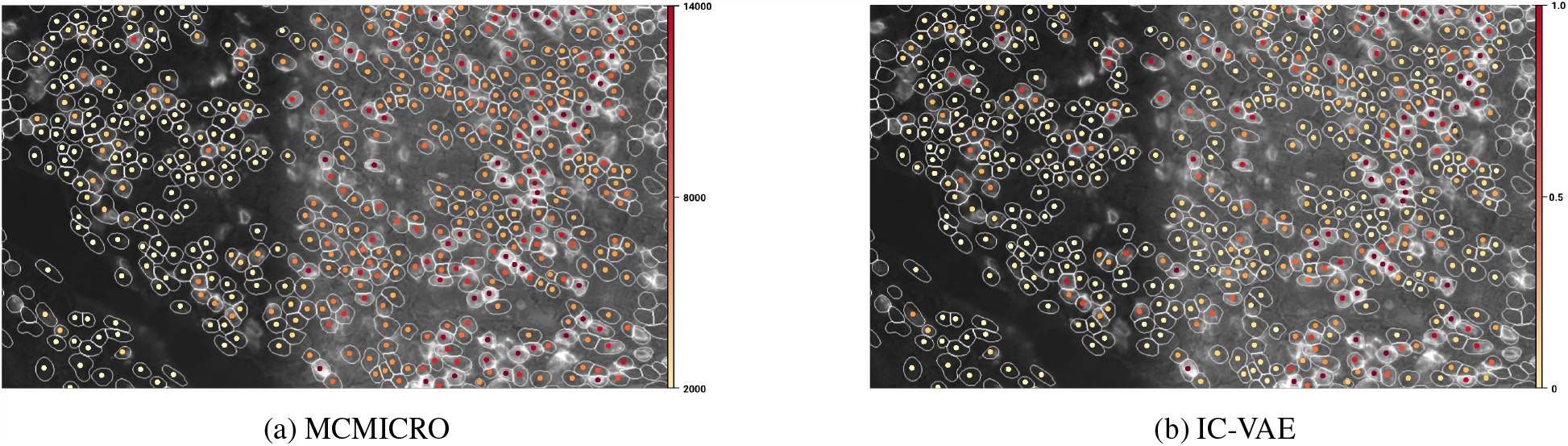
The *CD*3*D* expression quantification in the regions with and without background signal. (a) From the result of the MCMICRO, cells in the region with the high background signal always have a high expression value, regardless of whether they belong to the CD3+ or CD3-groups (b) The expression values from IC-VAE model are not affected bythe background signal.

**Figure 4:**
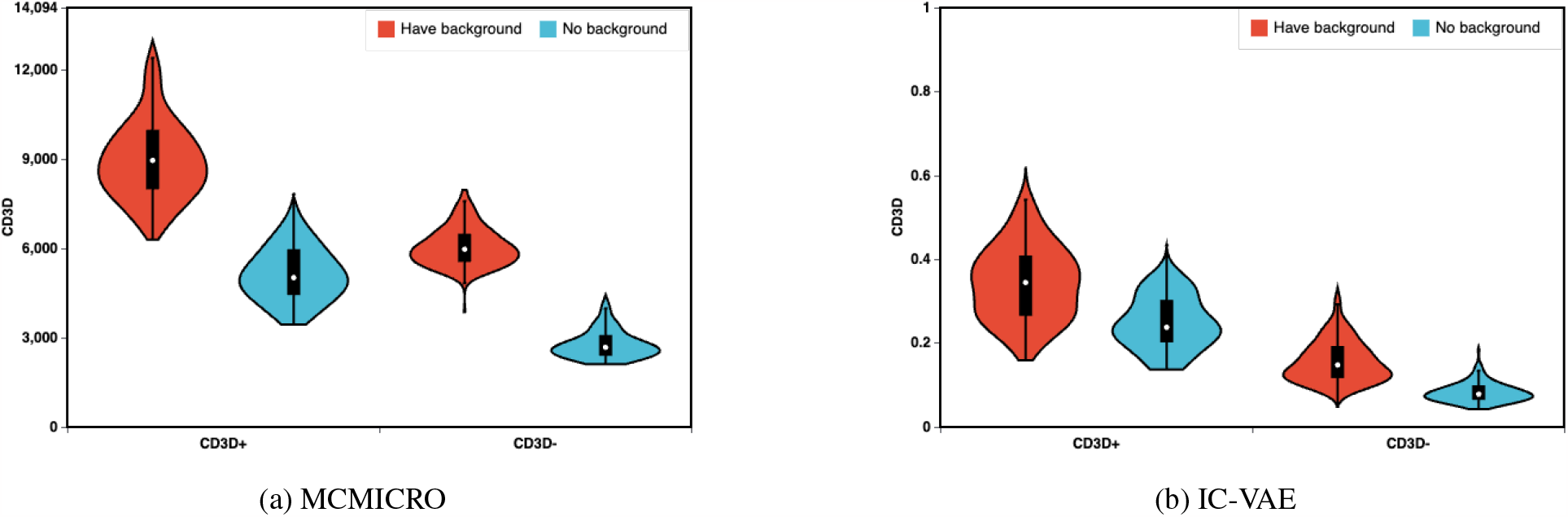
The protein expression distribution of *CD*3*D* between two groups CD3D+/CD3D-, divided by artifact region label. According to the result of MCMICRO, cells in the CD3-group located in regions with background signal even exhibit a higher level of protein expression compared to cells in the CD3+ group situated in regions without background signal

#### 3.3.2 Clustering cells for cell typing

In this experiment, we extract cells from three labeled regions and perform the clustering analysis as described in 3.2.5. Ideally, cells within a cluster should be the same cell type. We first evaluate clustering results using RandIndex [10] - given the cell-type annotation as the ground truth. As shown in Table 1, the RandIndex between the cell type label and IC-VAE clustering results are higher than the RandIndex between the cell type label and MCMICRO clustering results on all regions. This indicates that IC-VAE clustering results are more similar to the cell type annotations than MCMICRO clustering results.

**Table 1:**
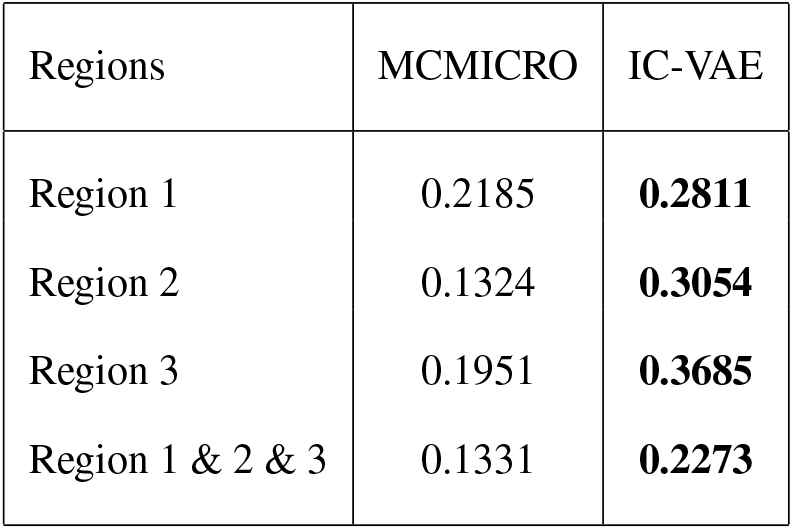
RandIndex between cell type labels and clustering result. IC-VAE outperforms MCMICRO in individual regions and in the combined region.

| Regions | MCMICRO | IC-VAE |
| --- | --- | --- |
| Region 1 | 0.2185 | **0.2811** |
| Region 2 | 0.1324 | **0.3054** |
| Region 3 | 0.1951 | **0.3685** |
| Region 1 & 2 & 3 | 0.1331 | **0.2273** |

We then further evaluate the clustering specificity to different cell type labels by:

- Assign all cells in a cluster with the label of the largest cell type population.
- For each cell type *A*, we create two binary vectors: the *truth*_*vector*, derived from the actual truth labels, and the *predict*_*vector* formed using the newly assigned labels. In these vectors, the *T RUE* value is assigned to cells labeled as *A*, and the rest are *FALSE*.
- Calculate the F1-score between each pair of *truth*_*vector* and *predict*_*vector*

As shown in Figure S3 and Table S2, the IC-VAE model has a higher specificity for cell type clustering. MCMICRO fails to separate the *epithelialcell* population from other cell types. Both methods fail to cluster the *fibroblast* and *regulatory T cell* populations out of the remaining cells.

#### 3.3.3 Visualize cell clusters on the UMAP

Similar to clustering, for the UMAP results, we expect cells from the same cell type label to be close together on the 2-D coordinates, while cells from different labels should be far apart. We evaluate the result for each cell type by:

- Normalize the UMAP coordinates to range from 0 to 1 for both x and y.
- Compute the centroid coordinates for each cell type based on the scaled UMAP coordinates of the cells within that cell type
- For each cell, determine its distance to the centroid of its respective cell type group, which is referred to as the *in_group*_*distance*. For *out_group*_*distance*, calculate the distance from the cell to the centroids of all other cell type groups
- The final score is defined as *median*(*out_group*_*distance*) / *median*(*in_group_distance*). The larger the score, the better the performance.

The results from Figure 5 and Table 2 indicate the better performance of the IC-VAE model. In the UMAP representation generated by IC-VAE, different cell-type clusters exhibit a greater degree of separation, with particular ease in distinguishing *myeloid leukocyte* and *lymphoma* clusters. In contrast, cells of the same cell type on the UMAP generated by MCMICRO form multiple clusters (*B cells* are split into 3 clusters, *myeloid leukocytes* are split into 2 clusters and *lymphomacells* are also split into 2 clusters). This observed phenomenon might be explained by the technical variance discussed in section 3.3.1. Furthermore, in the UMAP representation produced by MCMICRO, the *epithelial cell* population appears to be intermixed with a substantial portion of *myeloid leukocytes* and *B cells*. This outcome aligns with the clustering results from MCMICRO, where the epithelial cell population is also inseparable.

**Table 2:**
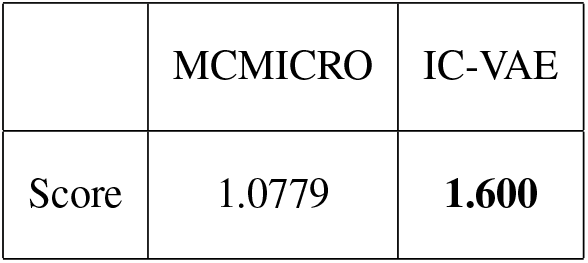
The ratio between *out_group_distance* and *in_group_distance* on the UMAP. If cells of the same cell type stay closer to each other, while cells of different cell types are more distinctly separated on the UMAP, the ratio will be higher

**Figure 5:**
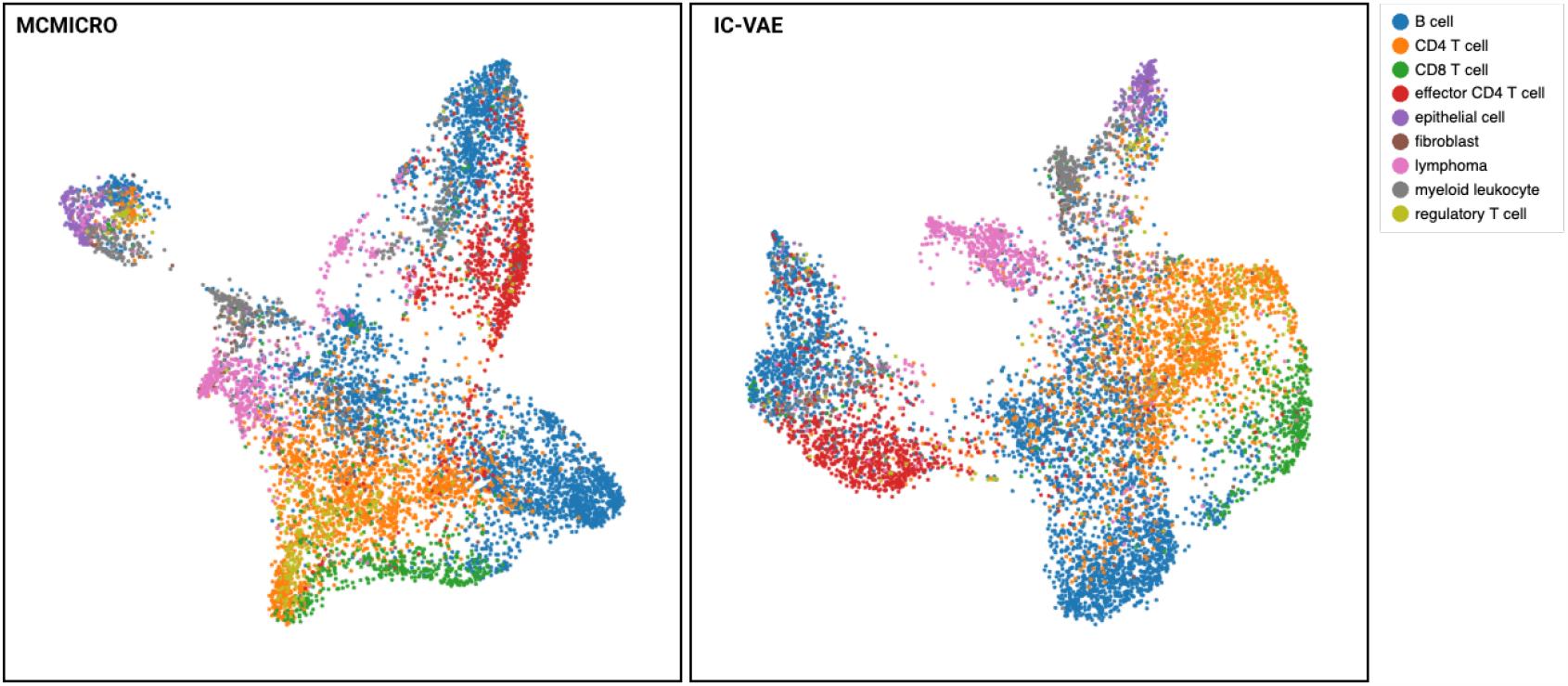
Comparative visualization of cell type distributions using UMAP. The UMAP plot from MCMICRO and IC-VAE with each color representing a different cell type. UMAP plot using IC-VAE is much more consistent with cell type annotations.

On both clustering and UMAP results, the IC-VAE model (and MCMICRO) fails to cluster the *regulatory T cell* and *fibroblast* population. To investigate, we visualize the expression matrix generated by IC-VAE (Figure S4). We found that the model had difficulty identifying the expression signal of *FOXP*3. Upon comparing this expression result with the original image, we observed that this was due to the fact that, unlike other marker expressions that exhibit distinct cellular patterns, *FOXP*3 expression was uniformly distributed within the cells (Figure 6. This uniform distribution closely resembled the background pattern, causing confusion for the model. These findings introduce new challenges that should be addressed in future works.

**Figure 6:**
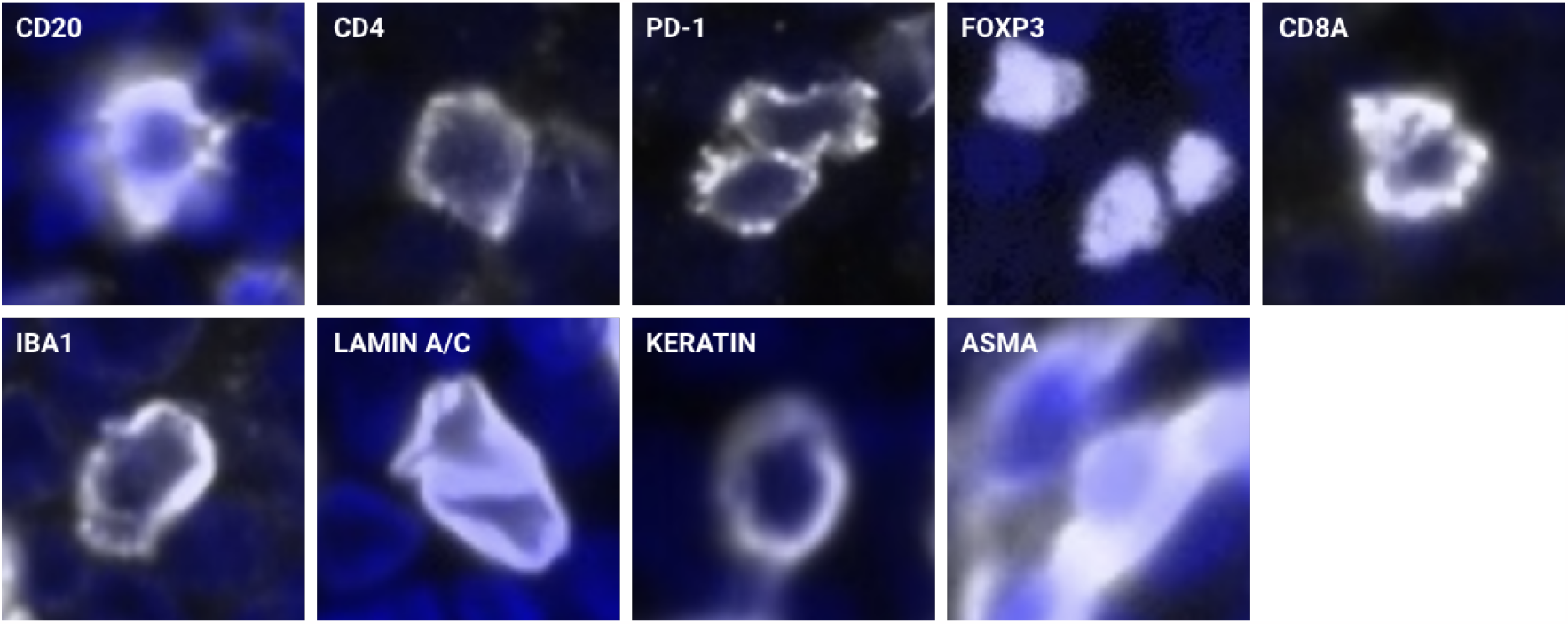
*FOXP*3 signal has a different pattern compared to other proteins, which leads to an incorrect estimation of *FOXP*3 expression in both methods.

## 4 Discussion

In this work, we introduce Information-Controlled Variational Autoencoder (IC-VAE). This leads to a significant improvement in interpreting protein expression from multiplexed tissue imaging data. The IC-VAE surpasses traditional approaches by allowing for controlled shared information among latent subspaces, which effectively separates true protein expression from cellular components and background noise. Unlike methods that rely on assumptions about cell symmetry and uniform protein distribution, IC-VAE accommodates the diverse cell shapes and intracellular structures resulting from 2-D tissue sectioning. Additionally, as our method just needs nucleus segmentation, it bypasses the need for adopting cell membrane segmentation, which usually provides inaccurate results. ^1^

The model provides more accurate protein quantification compared to the state-of-the-art tool MCMICRO, especially in densely packed cell regions where signals from neighboring cells often confound traditional analysis. We further showed that downstream applications, such as biomarker analysis, cell typing, and cell cluster visualization with t-SNE/UMAP, benefit from the precision of IC-VAE.

## Supplementary

### 1 MINE algorithm

#### Algorithm 1 MINE

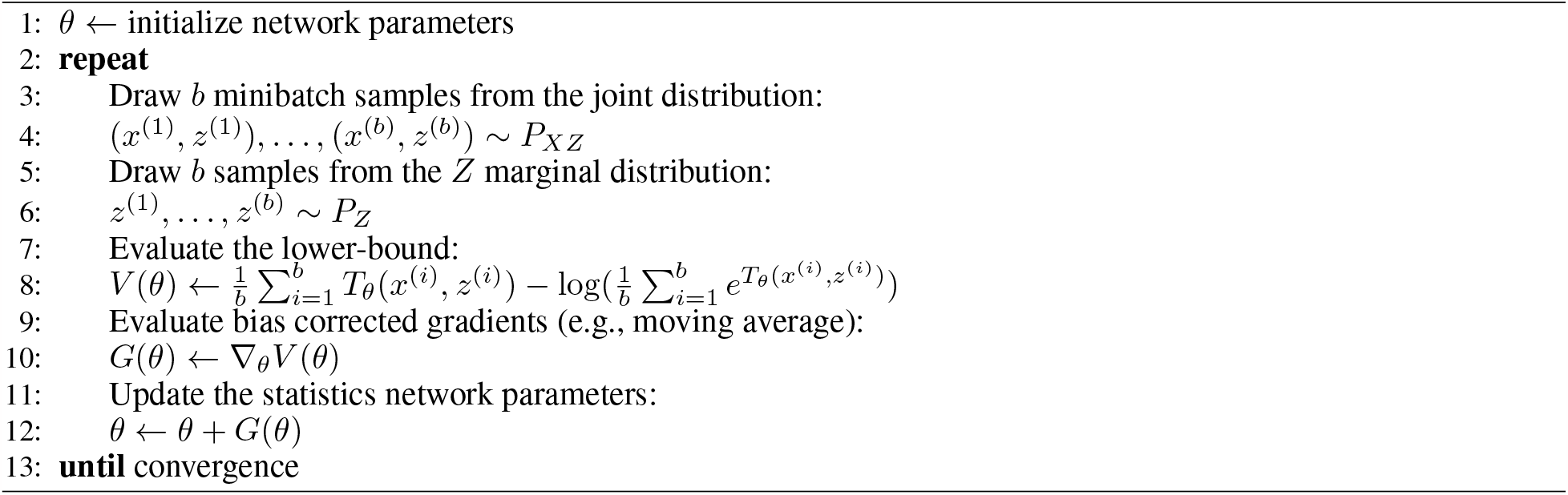

### 2 Cell type labeling

We annotate cell types based on the segmentation result and each individual image channel. From the list of input proteins, we define cell type using the markers listed in Table S1. The procedure for labeling each cell type is as below:

- Render the cell center onto the image, which is a combination of 1 DAPI channel and corresponding cell type biomarker channels (Figure S1)
- Select the cells that have the protein signal forming a complete cell structure (Figure S1a) and skip cells with no detectable signal (Figure S1b) or cells with noise/un-structured signal (Figure S1c)

The image channel for each biomarker across three selected region are shown in Figure S1.

### 3 Clustering evaluation result

### 4 Marker expression heatmap

**Table S1:**
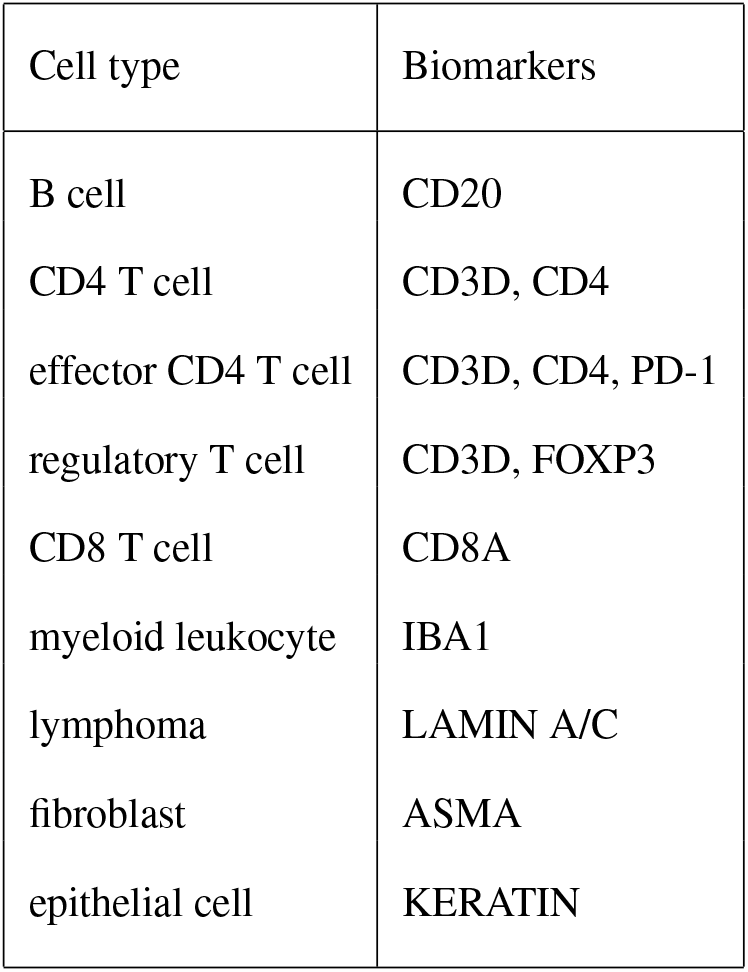
Biomarkers used for label each cell type.

| Cell type | Biomarkers |
| --- | --- |
| B cell | CD20 |
| CD4 T cell | CD3D, CD4 |
| effector CD4 T cell | CD3D, CD4, PD-1 |
| regulatory T cell | CD3D, FOXP3 |
| CD8 T cell | CD8A |
| myeloid leukocyte | IBA1 |
| lymphoma | LAMIN A/C |
| fibroblast | ASMA |
| epithelial cell | KERATIN |

**Figure S1:**
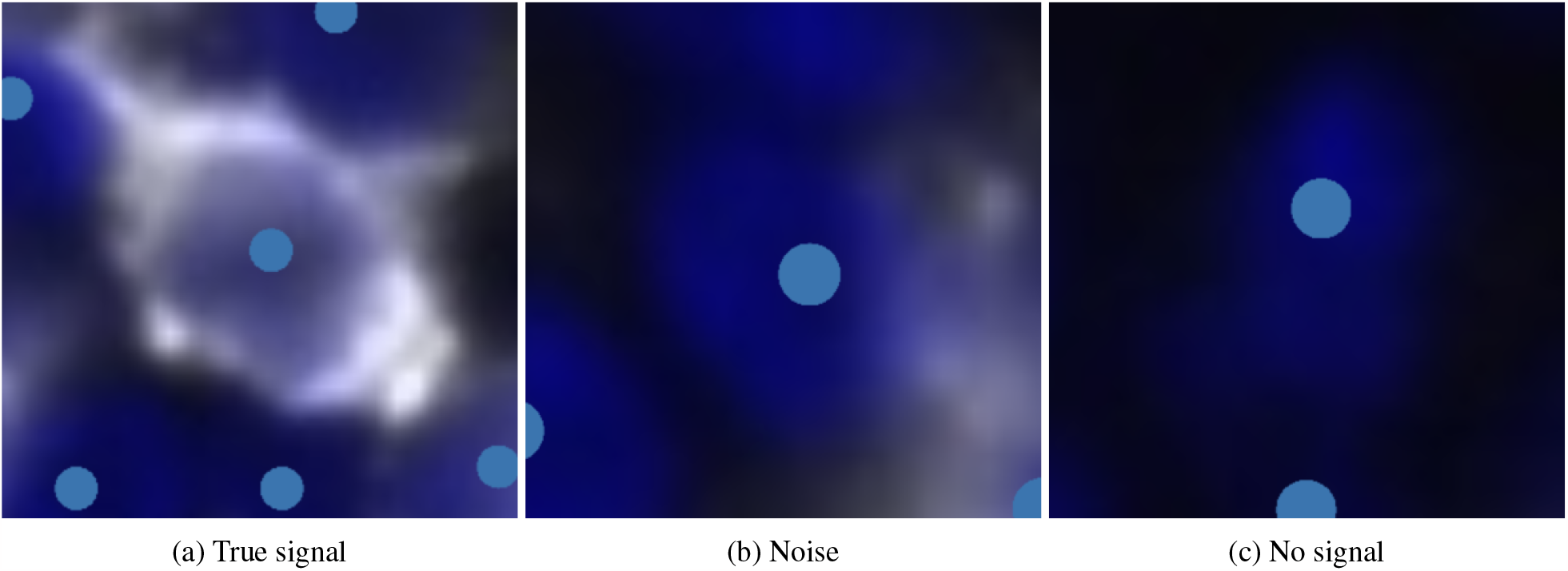
Distinguish between a cell’s true expression signal, background noise, and no signal. True cell type signal exhibits a consistent and complete cellular structure, which remains similar across cells of the same type. In contrast, noise signal displays a random pattern that varies from cell to cell and lacks the coherence seen in true cell type signal.

**Figure S2:**
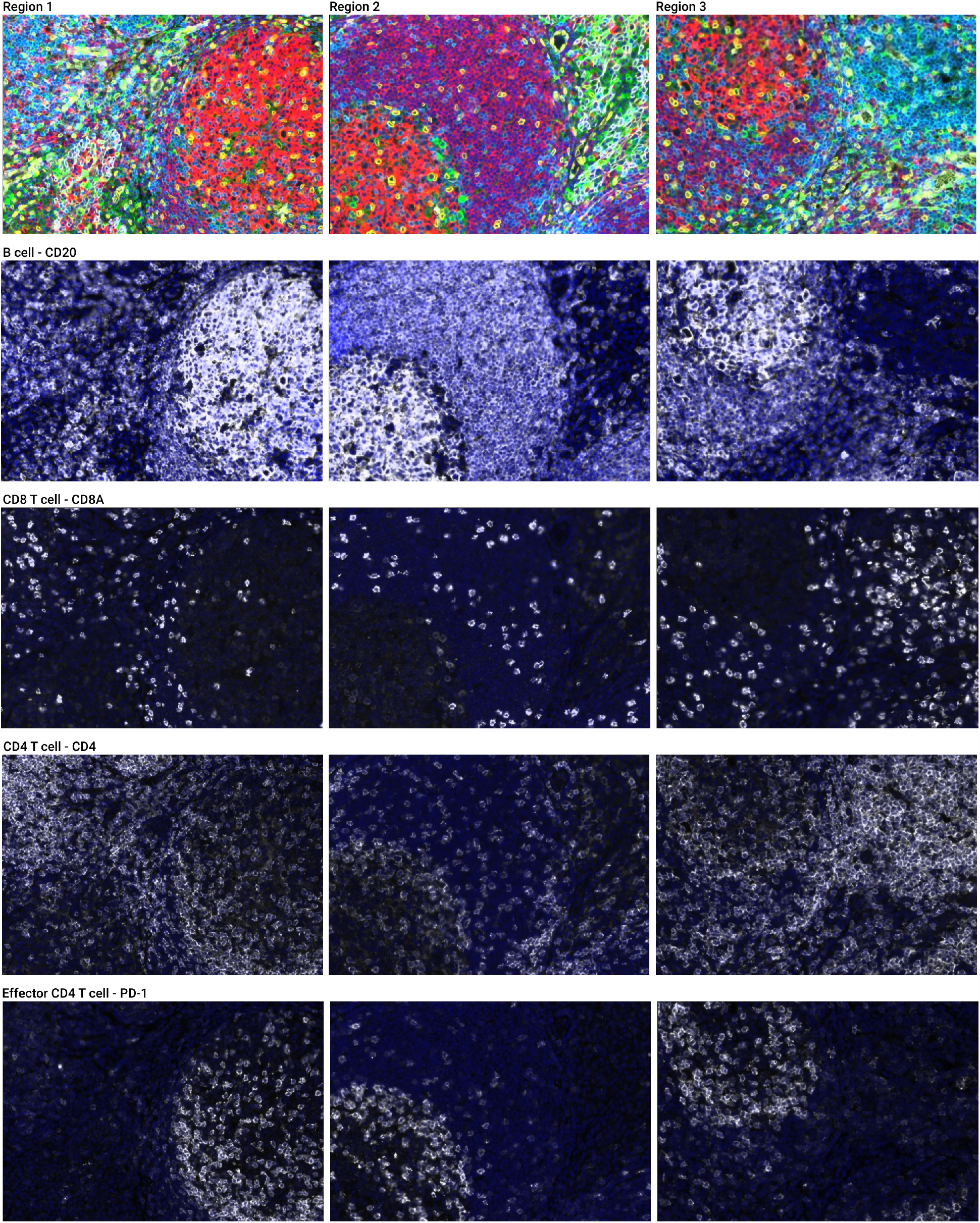
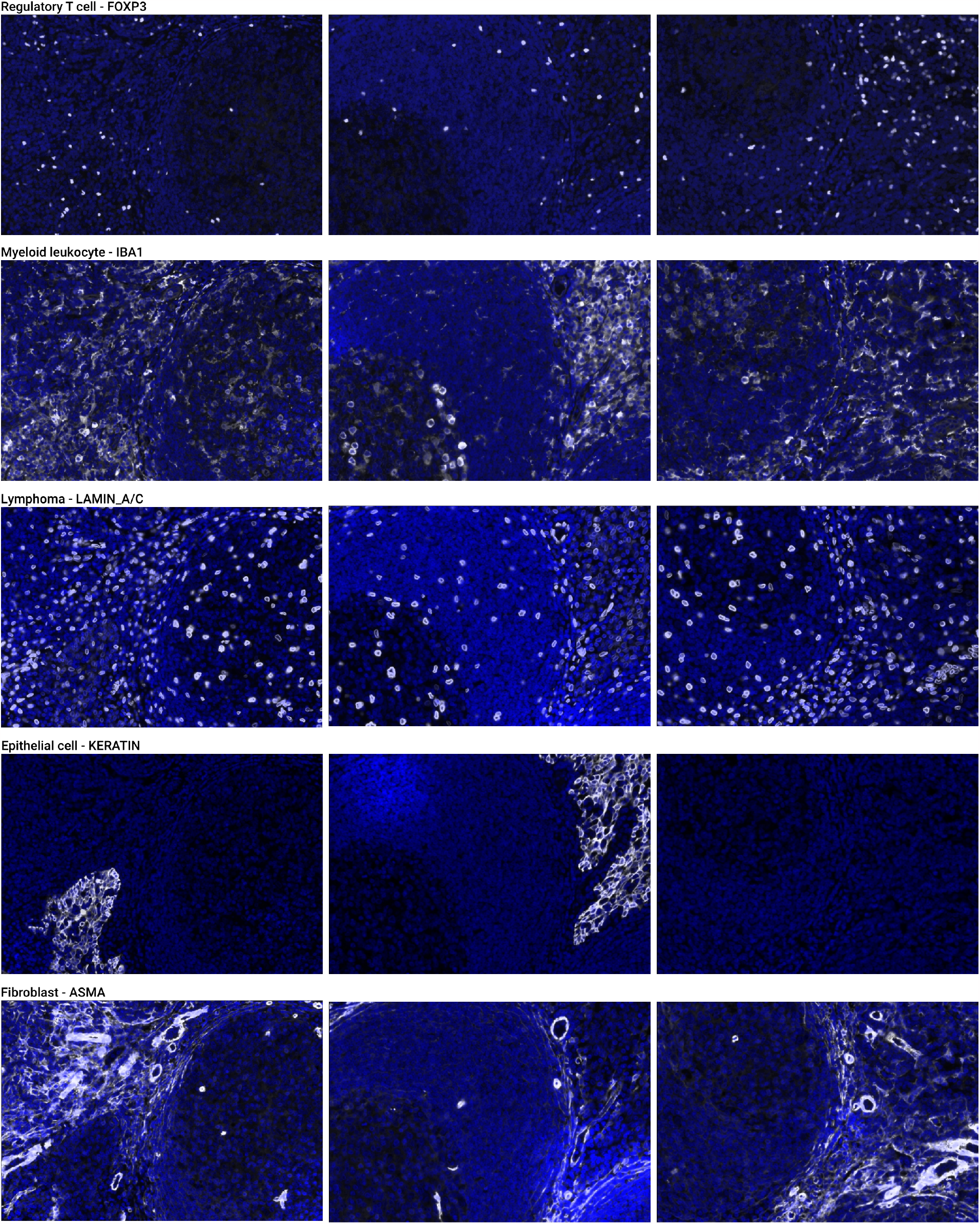
Protein channels used for manually labeling each cell type

**Figure S3:**
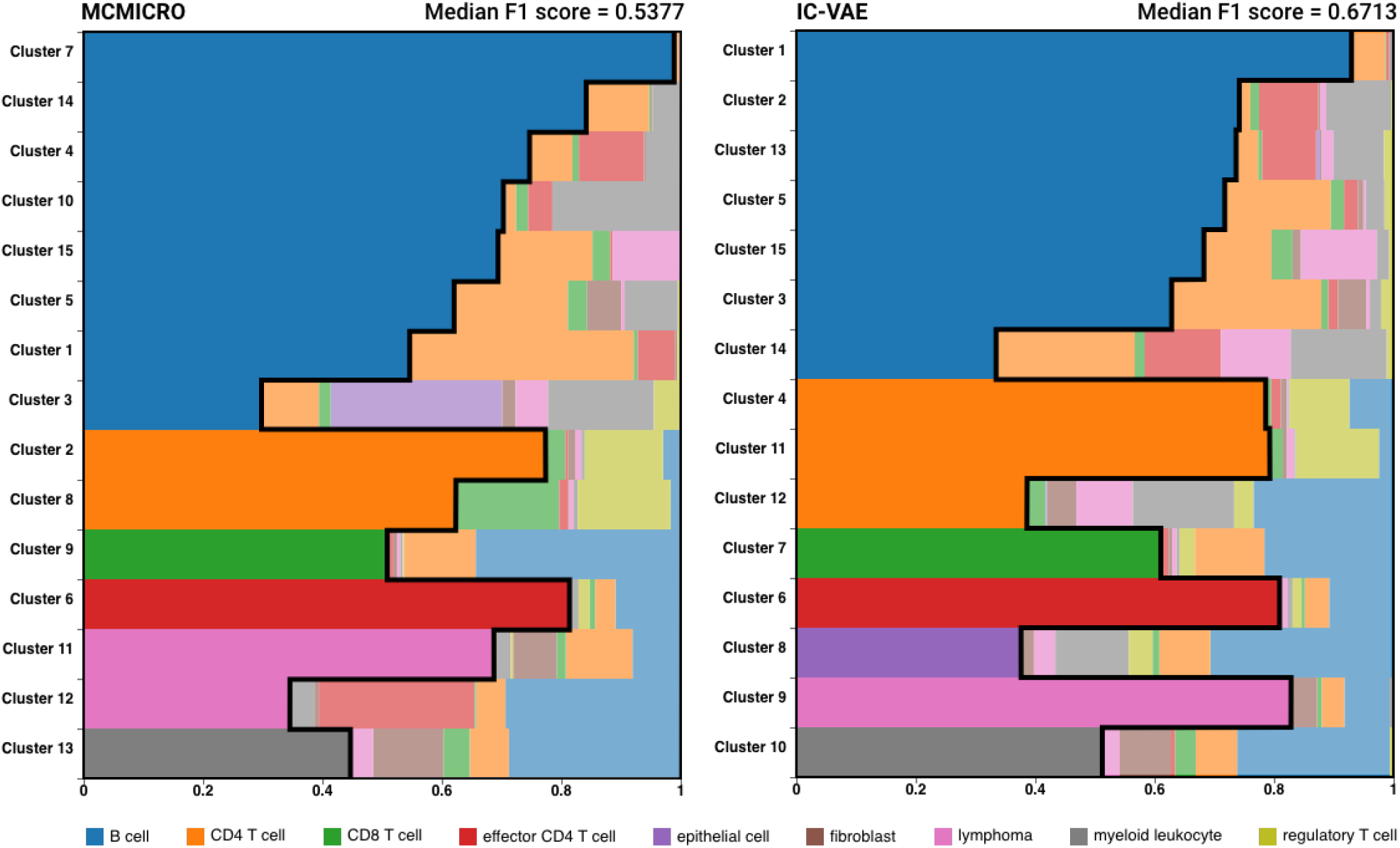
Composition plot between cell type label and clustering result of MCMICRO and IC-VAE. We then assign all cells from the cluster with the label of the largest cell type proportion. The epithelial cell population is missing from the MCMICRO result.

**Table S2:**
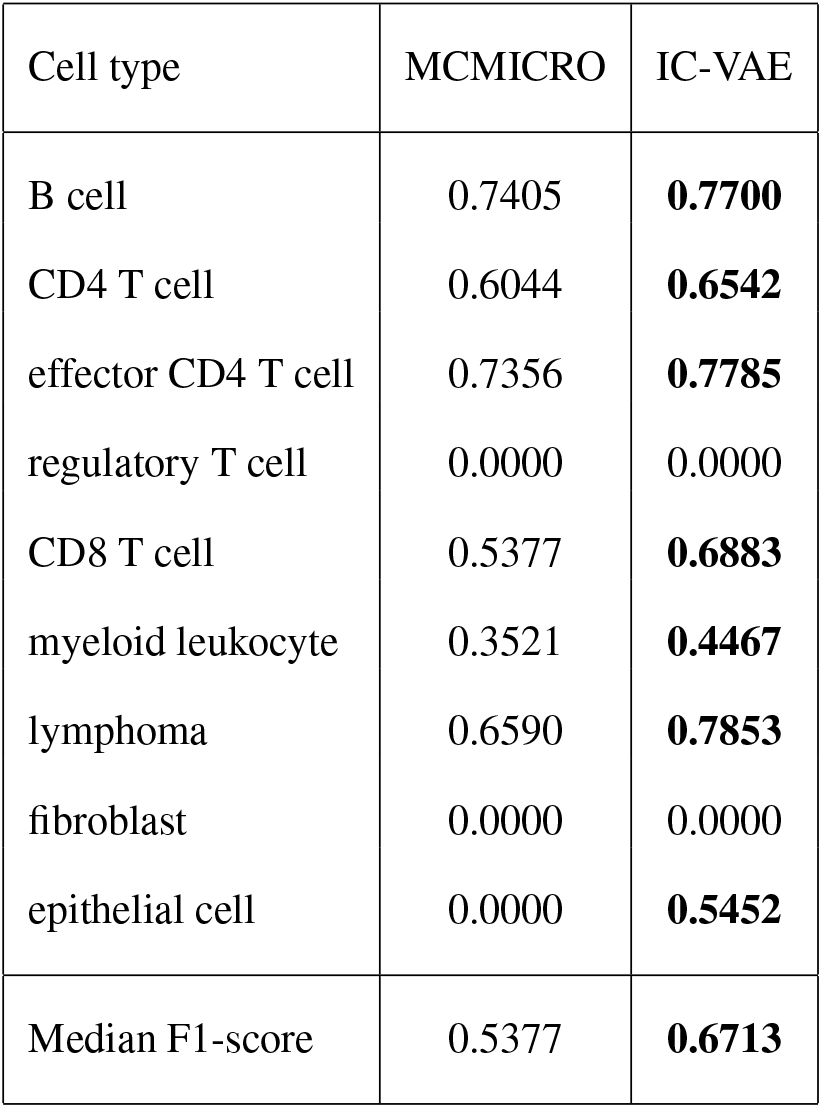
F1-score for each cell type between the truth label and the clustering assignment result.

| Cell type | MCMICRO | IC-VAE |
| --- | --- | --- |
| B cell | 0.7405 | **0.7700** |
| CD4 T cell | 0.6044 | **0.6542** |
| effector CD4 T cell | 0.7356 | **0.7785** |
| regulatory T cell | 0.0000 | 0.0000 |
| CD8 T cell | 0.5377 | **0.6883** |
| myeloid leukocyte | 0.3521 | **0.4467** |
| lymphoma | 0.6590 | **0.7853** |
| fibroblast | 0.0000 | 0.0000 |
| epithelial cell | 0.0000 | **0.5452** |
| Median F1-score | 0.5377 | **0.6713** |

**Figure S4:**
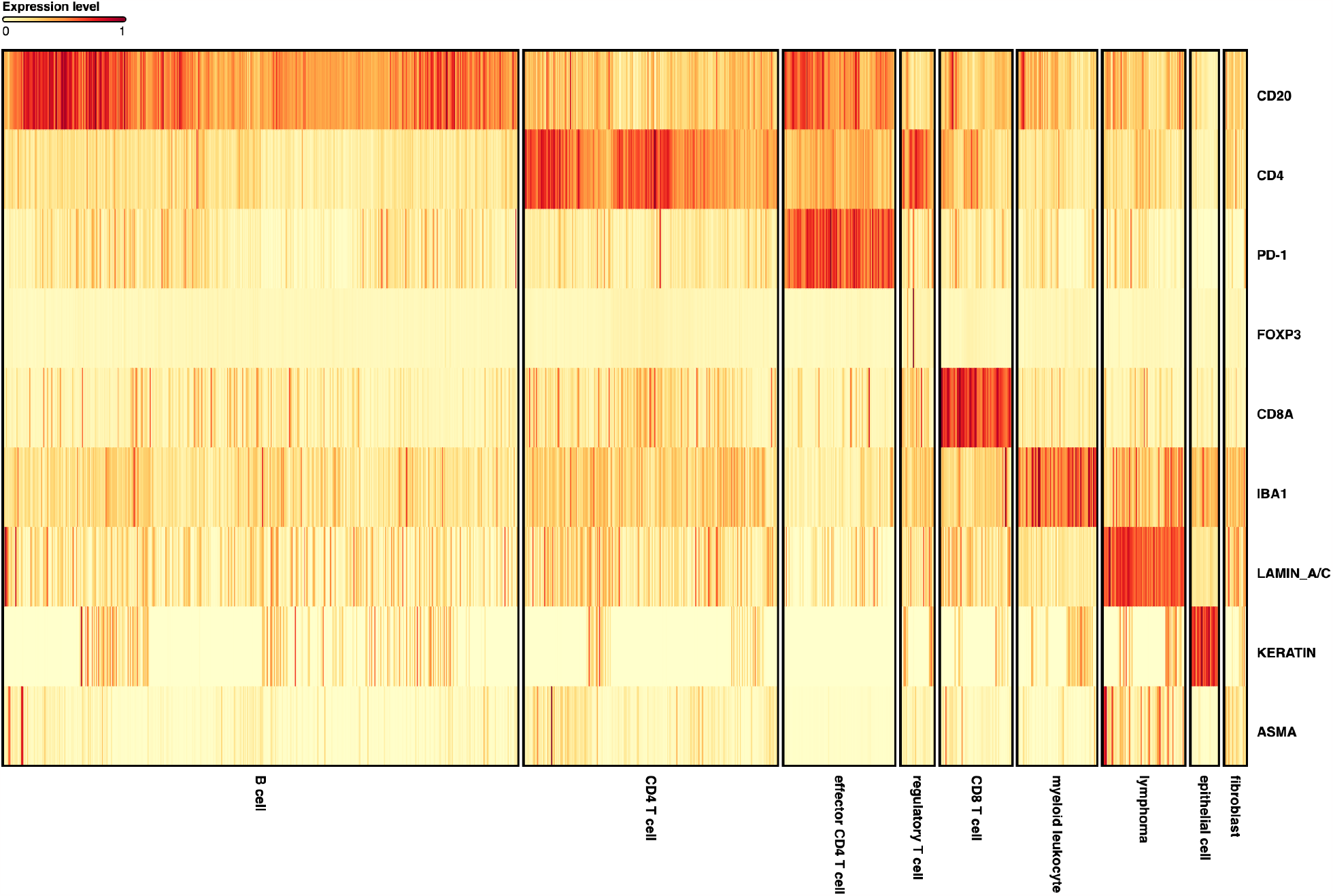
The heatmap of IC-VAE protein expression across cell type. FOXP3 shows no significant expression.

## Footnotes

1 Currently, many cell segmentation algorithms yield robust results for nucleus segmentation, but not for membrane segmentation.

## Notes

### Competing Interest Statement

Authors are working at BioTuring Inc., a Computational Biology company.

